# Chicory demonstrates substantial water uptake from below 2 m depth, but still did not escape topsoil drought

**DOI:** 10.1101/494906

**Authors:** Camilla Ruø Rasmussen, Kristian Thorup-Kristensen, Dorte Bodin Dresbøll

**Affiliations:** Department of Plant and Environmental Sciences, University of Copenhagen, Taastrup, Denmark

**Keywords:** *Cichorium intybus* L., Deep root growth, Deep water uptake, Drought resistance, Intercropping, hydrological tracer

## Abstract

**Aims:** Deep-rooted agricultural crops can potentially utilize deep water pools and thus reduce periods where growth is water limited. Chicory (*Cichorium intybus* L.) is known to be deep-rooted, but the contribution of deep roots to water uptake under well-watered and drought conditions by the deep root system has not been studied. The aim of this study was to investigate whether chicory could reach 3 m depth within a growing season and demonstrate significant water uptake from the deeper part of the root zone.

**Methods:** We tested if chicory exposed to either topsoil drought or resource competition from the shallow-rooted species ryegrass *(Lolium perenne L.)* and black medic (*Medicago lupulina L.)* would increase deep water uptake in compensation for reduced topsoil water uptake. We grew chicory in 4 m deep soil filled rhizotrons and found that the roots reached 3 m depth within a growing season.

**Results:** Water uptake from below 1.7 m depth in 2016 and 2.3 m depth in 2017 contributed significantly to chicory water use. However, neither drought nor intercropping increased the deep water uptake.

**Conclusion:** Chicory benefits from being deep-rooted during drought events, yet deep water uptake cannot compensate for the reduced topsoil water uptake during drought.

## Introduction

Minimizing water limitation during growth of agricultural crops is crucial to unlocking full yield potential. Crop yield losses vary according to the timing and severity of water limitations, but even short-term drought can be a major cause of yield losses (Zipper et al. 2016). Deep-rooted crops can potentially utilize otherwise inaccessible deep-water pools and thus reduce periods where crop production is water limited. In areas where precipitation is sufficient to rewet the soil profile during a wet season, shallow-rooted crops might still experience water limitation during the growing season, as they do not have access to the water stored deeper in the profile.

The potential influence of deep roots on water uptake has been highlighted numerous times (e.g. Canadell et al. 1996; Lynch and Wojciechowski 2015), still, information about the actual contribution of deep roots to water uptake remains scarce. Maeght et al. (2013) suggest that this is related to the absence of tools to measure deep root activity with sufficient throughput and standardization at affordable costs, and to the widespread assumption that as deep roots only represent a small fraction of the overall root system their contribution to root system function is marginal. It has also been questioned whether deep root growth under field conditions is too restricted by high soil strength, and unfavourable conditions such as e.g. hypoxia, acidity, and low nutrient availability, to substantially benefit the crop (Lynch and Wojciechowski 2015; Gao et al. 2016).

While some soils definitely restrict deep root growth, others have shown to allow roots to grow in the deeper soil layers (Sponchiado et al. 1989; Thorup-Kristensen and Rasmussen 2015). In addition, even though the majority of the root biomass is found in the topsoil, deep roots can contribute significantly to water supply in crops, as there is little connection between root biomass and root activity (Mazzacavallo and Kulmatiski 2015). Gregory et al. (1978) found that in the field, winter wheat had less than 5 % of its root biomass below 1 m depth, and as long as the water supply was sufficient in the upper meter, the biomass reflected the water uptake well. However, when the topsoil dried, the roots between 1 and 2 m depth supplied the plants with up to 20 % of the total water use. In a study conducted in an Amazonian tropical forest, Nepstad et al. (1994) found that they would have underestimated the evapotranspiration by 60 % in a dry year, had they not considered roots below 2 m depth.

Indirectly deeper root growth in crops has also been associated with deep-water uptake, as rooting depth has been shown to correlate positively with yield under drought in the field in e.g. wheat (Lopes and Reynolds 2010), bean (Sponchiado et al. 1989; Ho et al. 2005), rice (Uga et al. 2013) and maize (Zhu et al. 2010). In addition, modeling studies indicate that selection for deeper roots in grain crops could significantly improve deep-water acquisition and thereby yield in water deficit seasons (Manschadi A et al. 2006; Lilley and Kirkegaard 2011). Common to most of these studies is that deep root growth is considered to be in the range of 0.5 to 1.5 m depth. But several agricultural crops have the capability to grow roots below 2 m depth or even deeper within a season (Canadell et al. 1996; Ward et al. 2003; Thorup-Kristensen 2006; Rasmussen et al. 2015), and thereby get access to an extra pool of water originating from wet season surplus precipitation stored in the soil. For example, lucerne has shown to decrease the soil water content at 5 m depth (Fillery and Poulter 2006).

Hydrological isotope tracer techniques have over the last two decades become an increasingly popular tool to acquire information on temporal and spatial water use patterns in plants (e.g. Bishop and Dambrine 1995; Peñuelas and Filella 2003; Beyer et al. 2016). Injection of tracer into specific soil depths has proven to be a precise method to detect the relative water uptake from the chosen depth (Kulmatiski et al. 2010; Kulmatiski and Beard 2013; Bachmann et al. 2015; Beyer et al. 2016). The hydrological tracer techniques utilize the fact that no isotopic fractionation against isotope forms of hydrogen or oxygen occurs during soil water uptake by roots (Wershaw et al. 1966; Dawson and Ehleringer 1991; Bariac et al. 1994; Mensforth and Walker 1996).

The anthropocentric discussion of the importance of deep root growth in crop production is put in perspective by the fact that some plant species have evolved the potential to grow deep roots. Under what circumstances is that strategy beneficial? In this study, we hypothesize that deep root growth can help plants escape topsoil drought. More specifically, we aimed at testing the following hypotheses, using chicory *(Cichorium intybus L.)* as an example plant: 1) Chicory can grow roots below 3 m depth within a growing season. 2) Chicory has a significant water uptake from the deeper part of the root zone despite low root intensity. 3) When chicory is exposed to either topsoil drought or resource competition from shallow-rooted species, deep water uptake increases in compensation for the decreased topsoil water uptake.

Chicory is commonly grown in pasture mixtures for animal fodder or as a cash crop to produce inulin (Meijer et al. 1993). It is known to be able to reach at least 2.5 m depth (Thorup-Kristensen and Rasmussen 2015) and to be drought resistant (Monti et al. 2005; Skinner 2008; Vandoorne et al. 2012a). To test the hypotheses we grew chicory as a sole crop and in an intercropping with the two shallow-rooted species ryegrass *(Lolium perenne L.*) and black medic *(Medicago lupulina L.)* in 4 m deep rhizotrons. We allowed extensive root development before imposing a drought, as our focus was on the potential of deep roots to acquire water and not on deep root growth during drought.

## Materials and Methods

### Experimental facility

We conducted the experiment in a semi-field facility at University of Copenhagen, Taastrup (55°40’08.5”N 12°18’19.4”E), Denmark and repeated it for the two consecutive seasons, 2016 and 2017. We grew the crops in 4 m deep rhizotrons placed outside on a concrete foundation. The rhizotrons where 1.2 x 0.6 m rectangular columns constructed of steel frames. A waterproof plywood plate divided the rhizotrons lengthwise into an east- and a west-facing chamber with a surface area of 1.2 x 0.3 m. The rhizotrons stood on a north-south axis, narrow side facing towards one another (Fig. 1). On the east- and the west facing fronts of the rhizotrons, 20 transparent acrylic glass panels allowed inspection of root growth at the soil-rhizotron interface on the entire surface. Each panel was 1.2 m wide and could be removed to allow direct access to the soil column. Every third panel was 0.175 m tall, and the rest were 0.21 m tall. We used the narrow panels for placement of equipment and soil sampling. The tall panels were used only for root observations. To avoid disturbance of root growth, we never removed these panels during the experiment. All sides of the rhizotrons where covered in white plates of foamed PVC of 10 mm thickness to avoid light exposure of soil and roots. On the fronts, the foamed PVC plates were also divided into 20 panels. These were fixed in metal rails, allowing them to be slid off whenever we had to observe the roots. A wick in the bottom of the rhizotrons allowed water to drain out.

**Fig. 1.**
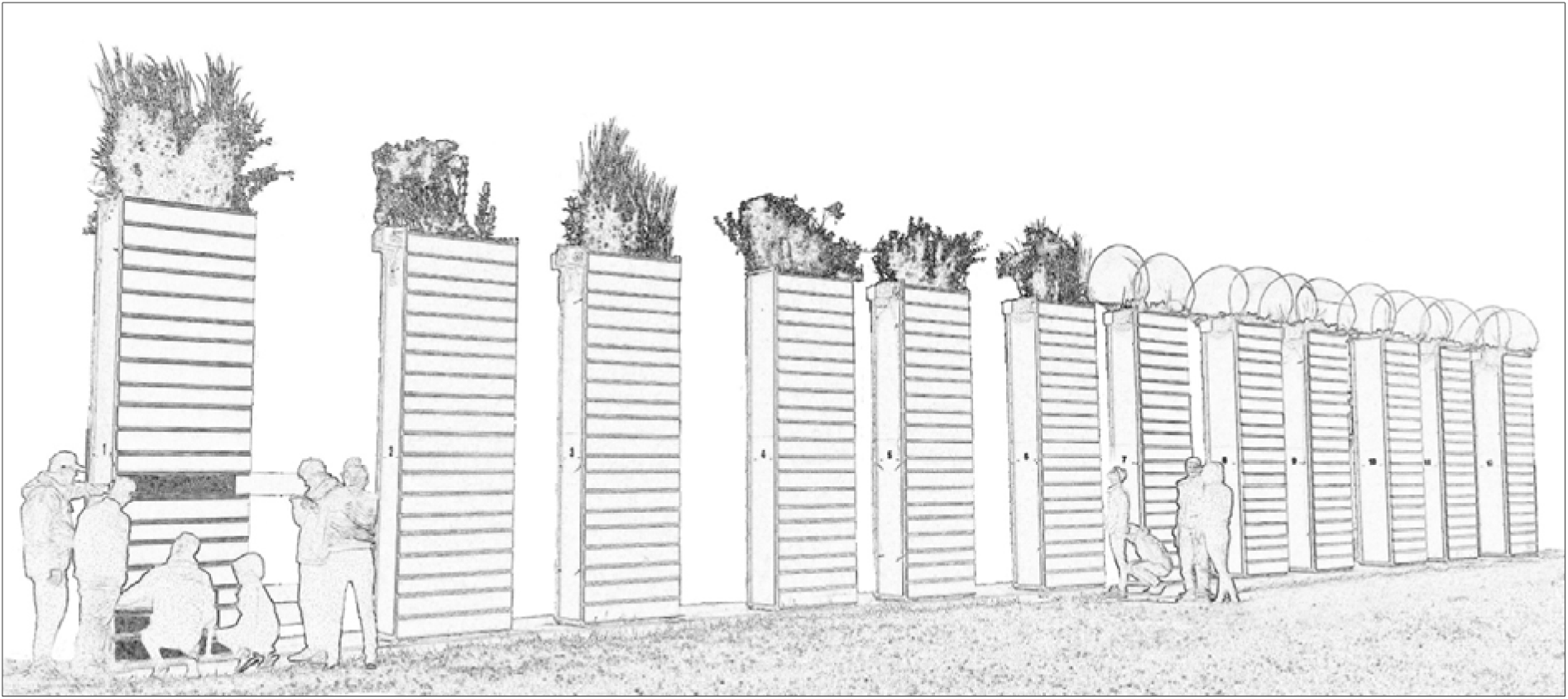
The rhizotron facility, consisting of 12 columns of 4 m height each divided into an east- and a west-facing chamber. See text for a detailed description

We used field soil as a growth medium. The bottom 3.75 m of the rhizotrons was filled with subsoil taken from below the plough layer at Store Havelse, Denmark (Table 1). We filled the upper 0.25 m with a topsoil mix of sandy loam and fine sandy soil, half of each, both from the University’s experimental farm in Taastrup, Denmark. To reach a soil bulk density comparable to field conditions we filled the soil into the rhizotrons stepwise at portions of approximately 0.15 m depth and used a steel piston to compact each portion by dropping it several times on the soil. We filled the rhizotrons in August 2015 and did not replace the soil during the two years. At the time of the experiment, average subsoil bulk density was 1.6 g m^-3^, which is close to field conditions for this soil type.

**Table 1:** Main characteristics of the soil used in the rhizotrons

| Depth (m) | Organic matter<sup>1</sup> (%) | Clay (%)<br><0.002 mm | Silt (%)<br>0.002-0.02 mm | Fine sand (%)<br>0.02-0.2 mm | Coarse sand (%)<br>0.2-2.0 mm | pH<sup>2</sup> |
| --- | --- | --- | --- | --- | --- | --- |
| 0.00-0.25 | 2.0 | 8.7 | 8.6 | 46.0 | 35.0 | 6.8 |
| 0.25-4.00 | 0.2 | 10.3 | 9.0 | 47.7 | 33.0 | 7.5 |

<sup>1</sup> Assuming that organic matter contains 58.7 % carbon.
<sup>2</sup> pH = Reaction Number (Rt) – 0.5. Measured in a 0.01 M CaCl<sub>2</sub> suspension, soil:suspension ratio 1:2.5.

We constructed rainout shelters to control water supply in the drought stress treatment. In 2016, we covered the soil with a transparent tarpaulin that had a hole for each plant stem. The tarpaulins were stretched out and fixed with a small inclination to let the water run off. It turned out that this design failed to keep out water during intense precipitation events, which happened twice during the season. Thus in 2017, we designed barrel roof rainout shelters instead, using the same clear tarpaulin and placed them on all rhizotrons. The rain-out shelters were open in the ends and on the sides to allow air circulation but were wider than the rhizotrons to minimize that water reached the chambers during precipitation under windy conditions.

We installed a drip irrigation pipe (UniRam™ HCNL) with a separate tap in each chamber. The pipe supplied 5 I hour^-1^, equivalent to 14 mm hour^-1^ according to the surface area of the growth chambers.

### Experimental design

We had two treatments in 2016 and four in 2017. In both years we grew chicory *(C. intybus L.,* 2016: cv Spadona from Hunsballe frø. 2017: cv Chicoree Zoom F1 from Kiepenkerl) in monoculture under well-watered (WW) and drought stress (DS) conditions. In 2017, we also grew chicory intercropped with either ryegrass (*L. perenne L.)* or black medic (*M. lupulina L*.), both in a WW treatment. For chicory, we chose to work with a hybrid vegetable type cultivar in the second year to reduce the variation among plants in size and development speed seen in the forage type used the first year. In 2016, we transplanted four chicory plants into each chamber. In 2017, we increased the number to six in order to reduce within-chamber variation. For the two intercropping treatments in 2017, we transplanted five plants of ryegrass or black medic in between the six chicory plants.

For the 2016 season, chicory plants were sown in May 2015 in small pots in the greenhouse and were transplanted into the rhizotrons 30 September. Despite our attempt to compact the soil inside the rhizotron chamber, precipitation made the soil settle around 10 % during the first winter. Therefore, 29 February 2016, we carefully dug up the chicory plants, removed the topsoil, filled in subsoil before filling topsoil back in and replanting the chicory plants. A few chicory plants did not survive the replanting and in March we replaced them with spare plants sown at the same time as the original ones and grown in smaller pots next to the rhizotrons. In 2017, we sowed chicory in pots in the greenhouse 11 April and transplanted them to the rhizotron chambers 3 May (Table 2). Chicory is perennial, it produces a rosette of leaves the first year and the second year it grows stems and flowers.

**Table 2:** Timeline of the experiments in 2016 and 2017

|  | 2016 | 2017 |
| --- | --- | --- |
| Sowing | May 2015 | 11 April |
| Transplanting | 29 February | 3 May |
| Onset of drying out | 26 June | 13 July |
| H<sup>2</sup> tracer study | 19-25 July | 15-21 August |
| Water uptake calculations | 24-27 July | 17-21 August |
| Harvest | 28 July | 11 September |

We grew all treatments in three randomized replicates. The soil inside the six chambers not used for the experiment in 2016 but included in 2017 had also sunken during the 2015/2016 winter and the same procedure was used to top up soil in these chambers before transplanting the chicory plants.

In 2016, we fertilized all chambers with NPK 5-1-4 fertilizer equivalent to 100 kg N ha^-1^, half 1 April and the other half 21 June. In 2017, we fertilized all chambers 3 May and 1 June following the same procedure. Two chambers were accidentally over irrigated mid-June 2017 and we refertilized them 16 June.

In 2016, we pruned the plants at 0.5 m height, several times between 24 May and 12 July to postpone flowering and induce leaf and root growth.

We started drying out the DS treatments 26 June in 2016 and 13 July in 2017, where we stopped irrigation and mounted the rainout shelters. In 2016, we kept irrigating the WW treatments whenever precipitation was considered insufficient to meet plant needs. In 2017, where the rainout shelters excluded precipitation in all chambers we kept irrigating all treatments apart from the DS to ensure sufficient water supply. However, we chose to supply the same amount of water in all the irrigated chambers, which led to different levels of soil water content.

### Biomass and^13^C

We harvested aboveground biomass 28 July in 2016 and 11 September in 2017. We dried the biomass at 80°C for 48 hours. The biomass was analysed for ^13^C/^12^C ratio on an elemental analyser interfaced to a continuous flow isotope ratio mass spectrometer (IRMS) at the University of California Stable Isotope Facility (Davis, California, USA). Isotope values are expressed in delta notation (δ) in per mill [‰] following the definition of Coplen (2011):

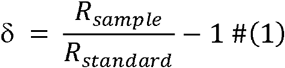

where *R_sample_* is the ratio of the less abundant to the more abundant isotope (^13^C/^12^C) in the PeeDee Belemnite (R_standard_ = 11180.2 × 10^~6^) was used. Analytical precision (σ) was 0.2‰.

The ^13^C/^12^C ratio in plants is directly related to the average stomatal conductance during growth, as discrimination between ^12^CO_2_ and ^13^CO_2_ during photosynthesis is greatest when stomatal conductance is high. When stomates are partially or completely closed, a greater part of the CO_2_ inside the leaf is absorbed resulting in less fractionation and thereby higher δ^13^C values of the plant tissue (Farquhar and Richards 1984; Farquhar et al. 1989).

### Root measurements

We documented the development in root growth by taking photos of the soil-rhizotron interface through the transparent acrylic glass panels. For this purpose, we designed a “photo box” that could be slid on the metal rails in place of the foamed PVC panels, and thereby excluded the sunlight from the photographed area. We placed a light source consisting of two bands of LED’s emitting light at 6000 K in the photo box. We used a compact camera (Olympus Tough TG 860). For each 1.2 m wide panel we took four photos to cover the full width of the panel. We photographed the roots 21 June and 18 July 2016 and 6 July, 16 August and 12 September 2017, corresponding to the time of drought initiation in the DS treatment, ^2^H tracer injection (see below) and for 2017, harvest. In 2017, harvest was postponed until 20 days after the ^2^H tracer-experiment, due to other tests running in the facility.

We recorded the roots using the line intersects method (Newman 1966) modified to grid lines (Marsh 1971; Tennant 1975) to calculate root intensity, which is the number of root intersections m-1 grid line in each panel (Thorup-Kristensen 2001). To make the counting process more effective we adjusted the grid size to the number of roots, i.e. we used coarser grids when more roots were present and finer grids for images with only a few roots. This is possible because root intensity is independent of the length of gridline. We used four grid sizes: 10, 20, 40 and 80 mm. To minimize the variance of sampled data we used grid sizes that resulted in at least 50 intersections per panel (Ytting 2015).

### Soil water content

We installed time-domain reflectometry sensors (TDR-315/TDR-315L, Acclima Inc., Meridian, Idaho) at two depths to measure volumetric water content (VWC) in the soil. In 2016, the sensors were installed at 0.5 and 1.7 m depth. In 2017, the sensors were installed at 0.5 and 2.3 m depth. Soil water content was recorded every 5 min in 2016 and every 10 min in 2017 on a datalogger (CR6, Campbell Scientific Inc, Logan, Utah). Discrepancies in measured VWC among the sensors at field capacity (FC) let us conclude that the sensors were precise but not particularly accurate, meaning that the change over time in VWC was reliable but not the measured actual VWC. We have therefore estimated a sensor reading for each sensor at FC and reported changes in VWC from FC. We estimated FC as the mean VWC over a 48-hour interval. In 2017, the measurement was made in the autumn after excess water from a heavy rainfall had drained away. In the autumn, there is little evaporation and no plant transpiration to decrease VWC below FC, making it an optimal time to estimate FC. We did not have data from autumn 2016, so instead, we estimated FC in early spring.

### Water uptake

We estimated water uptake from the VWC readings. We assume that water movement in the soil is negligible when VWC is below FC. Hence, the decrease in VWC can be interpreted as plant water uptake. Water uptake is therefore estimated as the mean decrease in VWC over a given time interval. We attempted to use intervals corresponding to the time of the ^2^H tracer studies. In 2016, the interval was a postponed a few days and in 2017, the time interval did not cover the first two days of the tracer study.

For the period from onset of drought to harvest 2017, we tested whether the daily water uptake at 2.3 m depth was affected by daily mean VWC at 0.5 m depth across all treatments. For this period, the VWC at 2.3 m depth was close to FC in all treatments and therefore unlikely to affect the water uptake. As transpiration demand is high at this time of the year and plants are large, we assumed that topsoil water limitations would limit total water uptake unless it is balanced by an increased water uptake lower in the profile. We excluded days in which the chambers were irrigated and one day after irrigation events to exclude periods with large soil water movement.

### ^2^H tracer

We used ^2^H labeled water injected into 2.3 m depth to trace water uptake from this depth. We mixed 90% ^2^H_2_O tracer with tap water 1:1, to achieve an enrichment of δ 5,665,651 ‰ and injected 100 ml per chamber. We removed one of the acrylic panels in each chamber temporarily to allow tracer injection and distributed it over 100 injection points in the soil. The injections were made at two horizontal rows of each 10 equally distributed holes 5 cm above and below 2.3 m depth respectively. In each of these 20 holes, we injected 5 ml tracer distributed between five points: 5, 10, 15, 20 and 25 cm from the horizontal soil surface. Tracer injection was made 19 July 2016 and 15 August 2017.

We captured the tracer signal by collecting transpiration water using plastic bags. For studies using tracers, collecting transpiration water is considered valid, as the tracers increase the enrichment level several orders of magnitudes, which make the fractionation negligible (Thorburn and Mensforth 1993; Beyer et al. 2016). We sampled the transpiration water one day before tracer injection as a control and one, two, three, four and six days after in 2016, and three and six days after in 2017. We fixed a plastic bag over each plant with an elastic cord that minimized air exchange with the surroundings. Transpiration water condensed on the inside of the plastic bag, which was folded inwards under the elastic cord to create a gutter for the water drops. Plastic bags were mounted on the plants two hours before noon and removed at noon.

We removed the plastic bags one by one, shook them to unite the drops, and transferred each sample to a closed plastic beaker. Later we filtered the samples through filter paper to remove soil and debris contamination and transferred the samples to glass vials.

We collected water from all plants and in most cases mixed the individual plant samples before analysis, taking equal amounts of water from each sample. Day 2 in 2016 and day 6 in 2017, we analysed the samples from each plant separately to get data on within chamber variation. For the control samples in 2017, we only collected water from two plants of each species per chamber. Single plant sample sizes varied from almost nothing to up to around 60 ml in 2016 and 30 ml in 2017. The amount did not only reflect differences in transpiration rate, as it was impossible to avoid spill when removing the plastic bags, and therefore we choose to use equal amounts of water from each plant. For the control samples where variation was small, this is of minor importance. The relatively large sample sizes for most samples limited the concerns of fractionation due to evaporation during filtering and sample transfer.

The samples were analysed for ^2^H at Centre for Isotope Research, University of Groningen, The Netherlands on a continuous flow isotope ratio mass spectrometer (IRMS, Isoprime 10) combined with a chromium reduction system (Europa PyrOH, Gehre et al. 1996). Isotope values are expressed in delta notation (δ) as given in equation 1. *R_sample_* is the ^2^H/^1^H ratio in the sample and *R_standard_* for δ^2^ H is Vienna standard mean ocean water (*R_standard_* ≈ 1/6412). Analytical precision (σ) was 0.7‰.

In order to identify whether tracer was present in a sample, we adapted the criteria proposed by Kulmatiski (2010). If a sample had a δ^2^ H-value at least two standard deviations higher than the control samples, tracer was assumed to be present.

### Statistics

Data analyses were conducted in R version 3.4.4 (R Core team 2018). The effect of treatment on aboveground biomass of chicory, black medic and ryegrass was tested in a mixed effects one-way ANOVA. Separate harvest of single plants allowed the inclusion of chamber as random effect to account for the fact that the two intercropped species are not independent.

The effect of soil depth and treatment on root intensity was tested in a mixed effects two-way ANOVA. We included chamber as random effect to account for the fact that the different depths are not independent. To meet assumptions of normality, depths where at least one of the treatments had no roots in any of the replicates, were excluded from the model. Separate analyses were made for each date.

The effect of soil depth and treatment on water uptake during a given time interval was tested in a mixed effect three-way ANCOVA with time as covariate. In 2016, we excluded the sensors from one replicate of the DS treatment because water reached it during a cloudburst. In 2017, we excluded two of the sensors at 0.5 m depth from the analysis, one in a chicory and ryegrass intercropping treatment and one in a chicory and black medic intercropping treatment. The first due to noise in the readings and the second due to readings showing a pattern in VWC that did not resemble the pattern of any of the other sensors.

The effect of treatment and time on ^2^H concentration in transpiration water was tested in a mixed effect two-way ANOVA. We log-transformed the response variable to meet the assumptions of homoscedasticity.

The effect of treatment on δ^13^C was tested in a one-way ANOVA. For 2017, the model is a mixed effects model because samples for each plant were analysed separately.

In all cases, separate analyses were made for each year. All models used met the assumptions of normality and homoscedasticity. Differences were considered significant at P <0.05. Tukey test P-values for pairwise comparisons were adjusted for multiplicity, by single step correction to control the family-wise error rate, using the multcomp package (Hothorn et al. 2008). For root intensity, we decided to control the family-wise error rate for each root depth. For the ^2^H concentration, we only made pairwise comparisons for the last date.

## Results

Plants grew well both years, and as hypothesized, roots were observed below 3 m depth by the end of the growing season. Both the uptake of ^2^H tracer and sensor readings showed that chicory acquired water from 2.3 m depth. However, our results do not suggest that compensation takes place, i.e. deep water uptake was not increased to balance the decreased topsoil water uptake during drought.

### Biomass

Plant development differed between the two experimental years. In 2016, the chicory plants were in their second growth year and went into the generative stage right from the start of the growing season. They started flowering in late May. In 2017, the chicory plants were in their first year of growth and stayed in the vegetative state. Aboveground biomass of chicory did not differ significantly between the two treatments in 2016 and was 6.52 and 6.85 t ha ^-1^ in the WW and DS treatment, respectively. In 2017, chicory biomass was 4.65 and 3.64 ton ha^-1^ in the WW and DS treatment respectively and 2.80 and 2.211 ha^-1^ when intercropped with either black medic or ryegrass. Biomass of black medic and ryegrass was 5.89 and 7.68 ton ha^-1^ respectively. Both intercropping treatments significantly reduced chicory biomass compared to the WW treatment. Ryegrass produced significantly more biomass than black medic (Fig. 2).

**Fig. 2.**
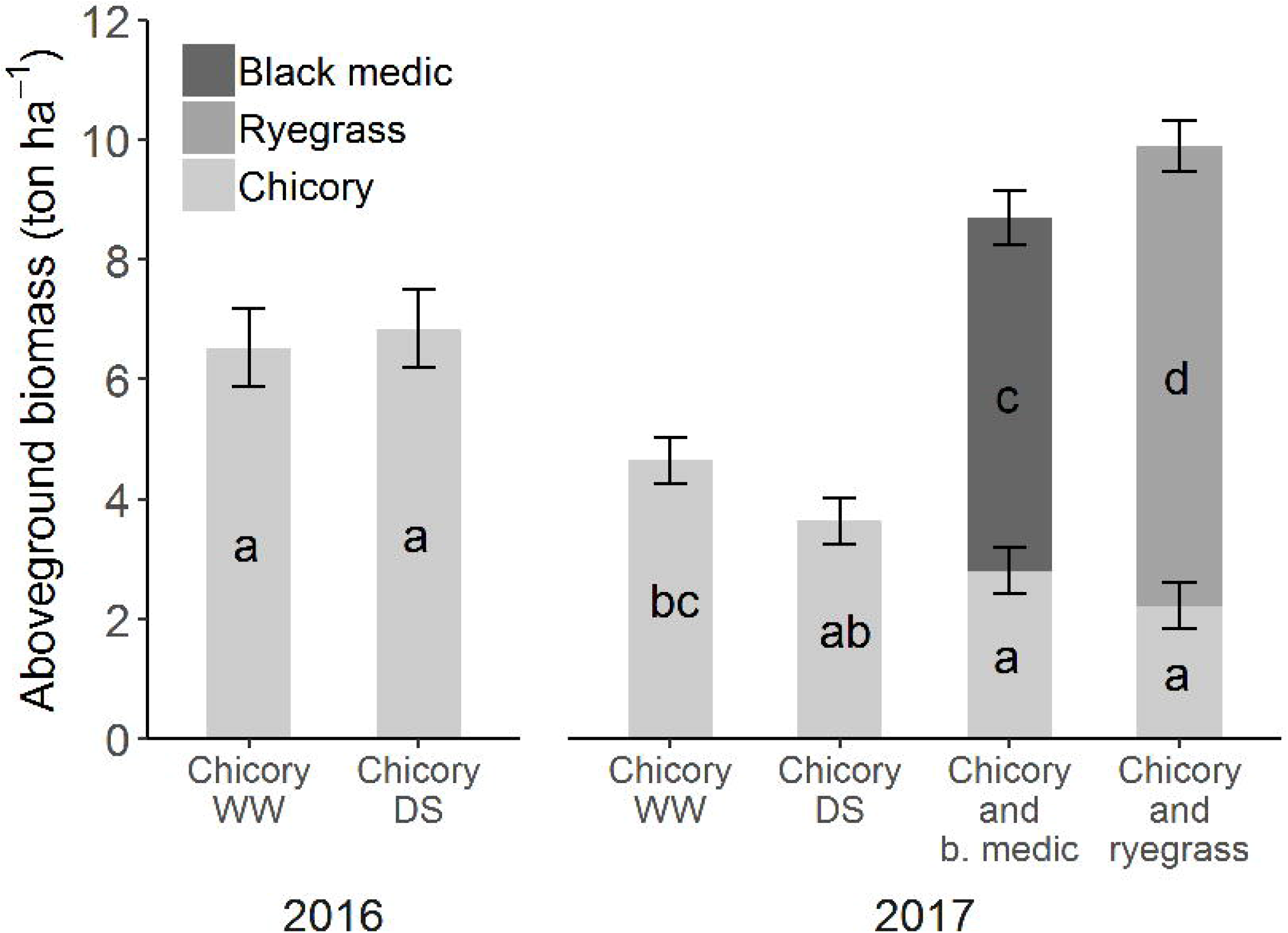
Biomass harvested 28 July 2016 and 11 September 2017 in the well-watered (WW) and drought stressed (DS) chicory sole crop treatments, and the chicory intercropping treatments with ryegrass and black medic respectively. Error bars denote standard errors, and letters indicate significant differences among treatments for each year. Part of the data has also been published in (Rasmussen et al. 2018).

### Root growth

Root growth showed a similar pattern across the four treatments; however intercropping decreased total root intensity down to around 2 m depth (only significant in few depths), except for 0.11 m depth, where the chicory and ryegrass intercropping treatment had a significantly higher root intensity than the other treatments. Roots of intercropped species could not be distinguished and the reported root intensities are thus the sum of two species in the intercropping treatments. The month-long summer drought did not influence root intensity in any depths.

In 2016, roots had reached 2 m depth at the time of drought initiation, which was 3.5 months after transplanting. A month later, at the time of tracer injection the rooting depth of chicory had increased below 3 m depth (Fig. 3a and b). In 2017, roots were observed almost to the bottom of the rhizotrons already at drought initiation, 2 months after transplanting. However, only a few roots were present below 2 m depth. At the time of tracer injection, which was again 3.5 months after transplanting root intensity had started to increase down to 2.5 m depth, and at harvest, 4.5 months after transplanting this was the case down to around 3 m depth (Fig. 3c, d, and e).

**Fig. 3.**
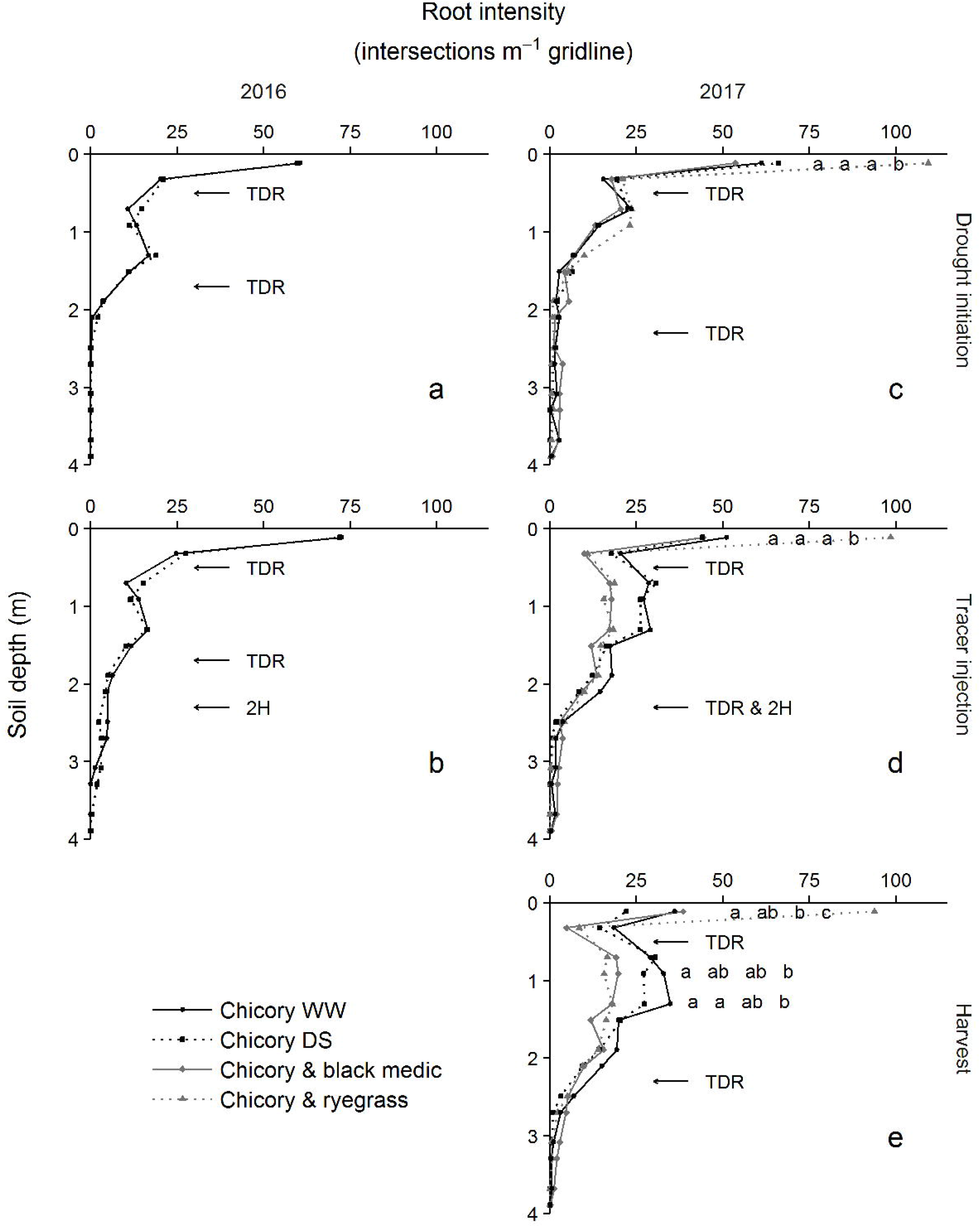
Root intensity in the well-watered (WW) and drought stressed (DS) chicory sole crop treatments and in the chicory intercropping treatment with ryegrass and black medic respectively in (a) 21 June 2016, (a) 18 July 2016, (c) 6 July 2017, (d) 16 August 2017 and (e) 12 September 2017, corresponding to the time of drought initiation in the DS treatment, ^2^H tracer injection and for 2017, harvest. Letters indicate significant differences among treatments in the given depth. Arrows indicate the depth of TDR sensors and ^2^H tracer injection.

### Soil moisture and water uptake

During the drought, 135 and 97 mm of water were excluded from the DS treatment in 2016 and 2017, respectively compared to the other treatments. In 2016, the soil dried out gradually at both 0.5 and 1.7 m depth in the DS treatment and in the WW treatment between the precipitation and irrigation events. As a result, the soil was drier in the DS than in the WW treatment at both depths recorded at the time of the tracer-experiment (Fig. 4a and b).

**Fig. 4.**
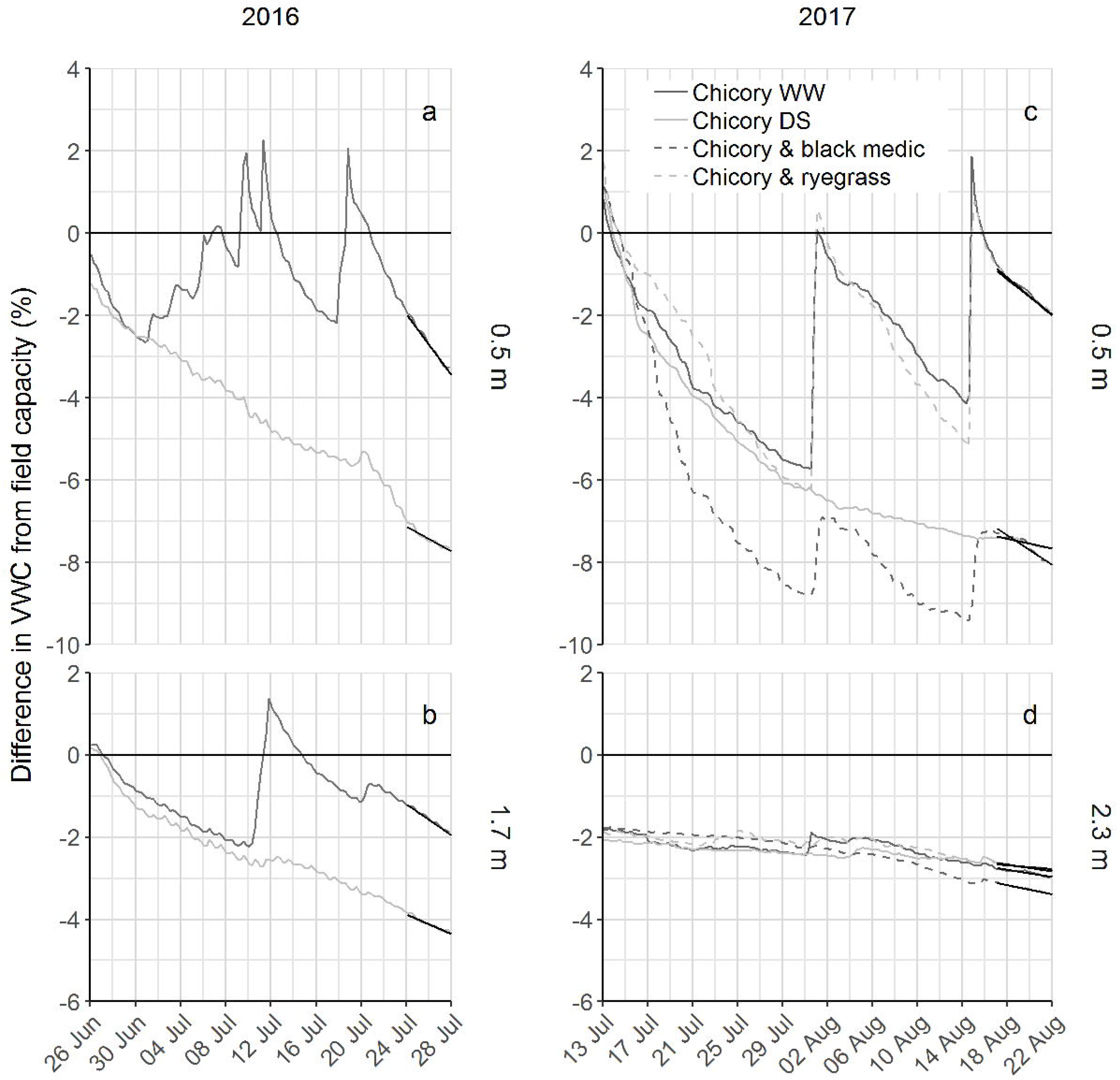
Difference in soil volumetric water content from field capacity at 0.5 and 1.7 m depth in 2016 and 0.5 and 2.3 m depth in 2017 in the well-watered (WW) and drought stressed (DS) chicory sole crop treatments and in the chicory intercropping treatment with ryegrass and black medic respectively. Line segments represent the outcome of a three-way ANCOVA on the time interval from 24 to 27 July in 2016 and 17 to 21 August in 2017. The slope of the segments gives the daily decrease in volumetric water content and is interpreted as daily plant water uptake. See also Fig. 5

Although chicory WW and the two intercropping treatments in 2017 received the same amount of water, less water reached the sensors at 0.5 m in the chicory and black medic intercropping than in the WW and the chicory and ryegrass intercropping. This indicates that the soil above the sensors was drier and therefore could withhold more water compared to the two other irrigated treatments. At the time of the tracer-experiment, soil water content under the chicory and black medic intercropping was similar to the DS treatment, which was lower in comparison to two other treatments (Fig. 4c and d).

During the tracer-experiment, chicory plants in the WW treatment acquired 3.7 and 2.3 mm water m^-1^ soil column day^-1^ from 0.5 m in 2016 and 2017, respectively. The uptake from 0.5 m depth was reduced by more than 50 % in the DS treatment compared to the WW treatment in both years. In the WW treatment, chicory took up 1.9 mm water m^-1^ soil column day^-1^ from 1.7 m depth in 2016, whereas the uptake was 0.44 mm water m^-1^ soil column day^-1^ from 2.3 m depth in 2017. In 2016, drought significantly reduced water uptake of chicory from 1.7 m depth, whereas no effect of drought was observed at 2.3 m depth in 2017. Common for both years was that the amount of water taken up from 0.5 m depth in the DS treatment was equal to the uptake from 1.7 and 2.3 m depth in 2016 and 2017 respectively. Both intercropping treatments significantly reduced water uptake at 0.5 m depth compared to the WW treatment, but no effect was seen at 2.3 m depth (Fig. 5).

**Fig. 5.**
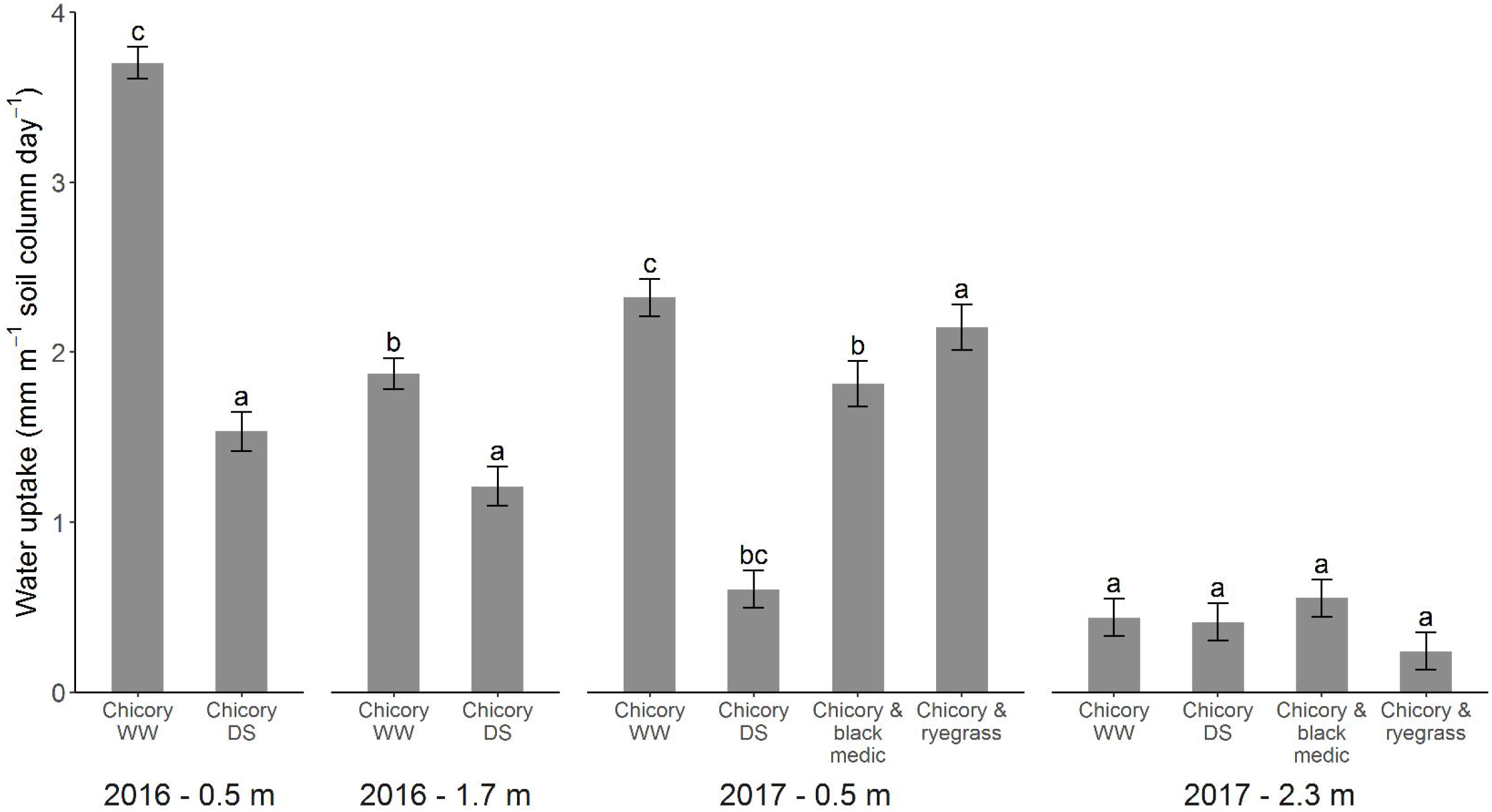
Mean daily decrease in soil volumetric water content at 0.5 and 1.7 m depth 24 to 28 July 2016 and 0.5 and 2.3 m depth 17 to 21 August 2017 in the well-watered (WW) and drought stressed (DS) chicory sole crop treatments and in the chicory intercropping treatment with ryegrass and black medic respectively. All days included. The daily decrease in volumetric water content is interpreted as daily plant water uptake. Error bars denote standard errors, and letters indicate significant differences among treatments in a three-way ANCOVA, with depth and treatment as factors and time as covariate. Separate analyses were made for each year

We did not find any effect of mean daily soil VWC at 0.5 m depth on water uptake at 2.3 m depth, giving no indication of compensatory deep water uptake (Data not shown).

### ^2^H enrichment

Chicory took up ^2^H tracer from 2.3 m depth in both years (Fig. 6a). Two days after tracer application in 2016, 21 out of 23 chicory plants demonstrated isotope ratios that were two standard deviations or more above controls. Six days after tracer application in 2017, it was 30 out of 64 chicory plants that showed the enrichment. No ryegrass or black medic plants indicated tracer uptake (Fig. 6b).

**Fig. 6.**
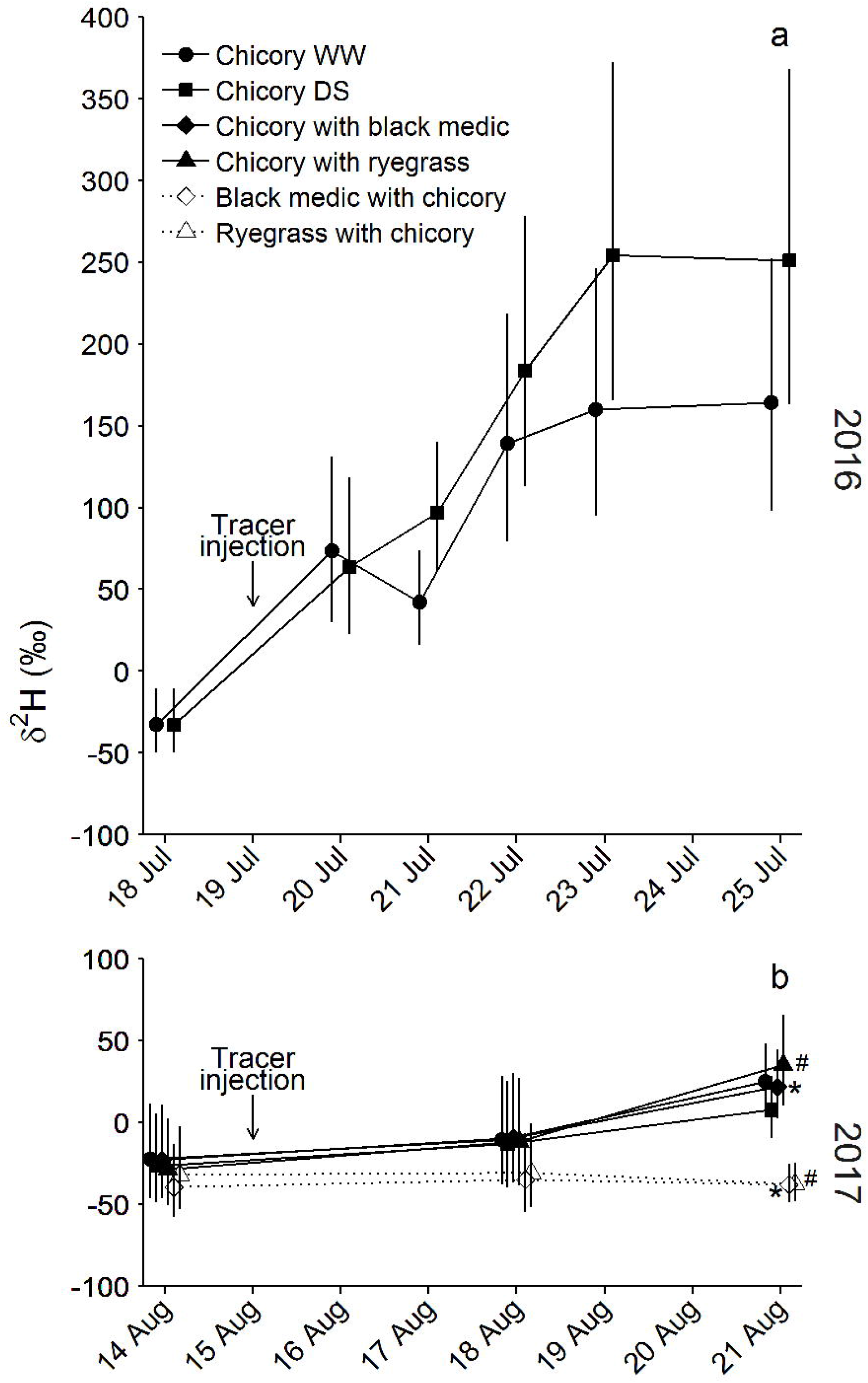
^2^H concentration in transpiration water before and after application of tracer at 2.3 m depth in (A) 2016 and (B) 2017 in the well-watered (WW) and drought stressed (DS) chicory sole crop treatments and in the chicory intercropping treatment with ryegrass and black medic respectively. We tested significant differences in a mixed effects two-way ANOVA. To meet the assumptions of homoscedasticity data were log-transformed. Separate analyses were made for each year and pairwise comparisons were only made for the last date. There was no effect of treatment in 2016. In 2017, the ^2^H concentration in chicory and in black medic in the intercropping treatment differed. Likewise in the chicory and ryegrass intercropping. Differing treatments are marked with identical symbols

In 2016, the ^2^H concentration in chicory plants in the DS treatment tended to be higher compared with the WW treatment, but the difference was not significant. In 2017, no differences were seen in tracer concentration among chicory plants across the treatments. Black medic and ryegrass plants revealed significantly lower ^2^H enrichment in comparison to intercropped chicory.

### δ^13^C enrichment

In 2016, there was no effect of drought on the ^13^C concentration of the chicory biomass (Fig. 7). Similarly, there was neither an effect of drought nor intercropping with ryegrass in 2017. However, intercropping with black medic increased the ^13^C concentration in chicory indicating that chicory was more drought stressed in this treatment than in any of the other treatments.

**Fig. 7.**
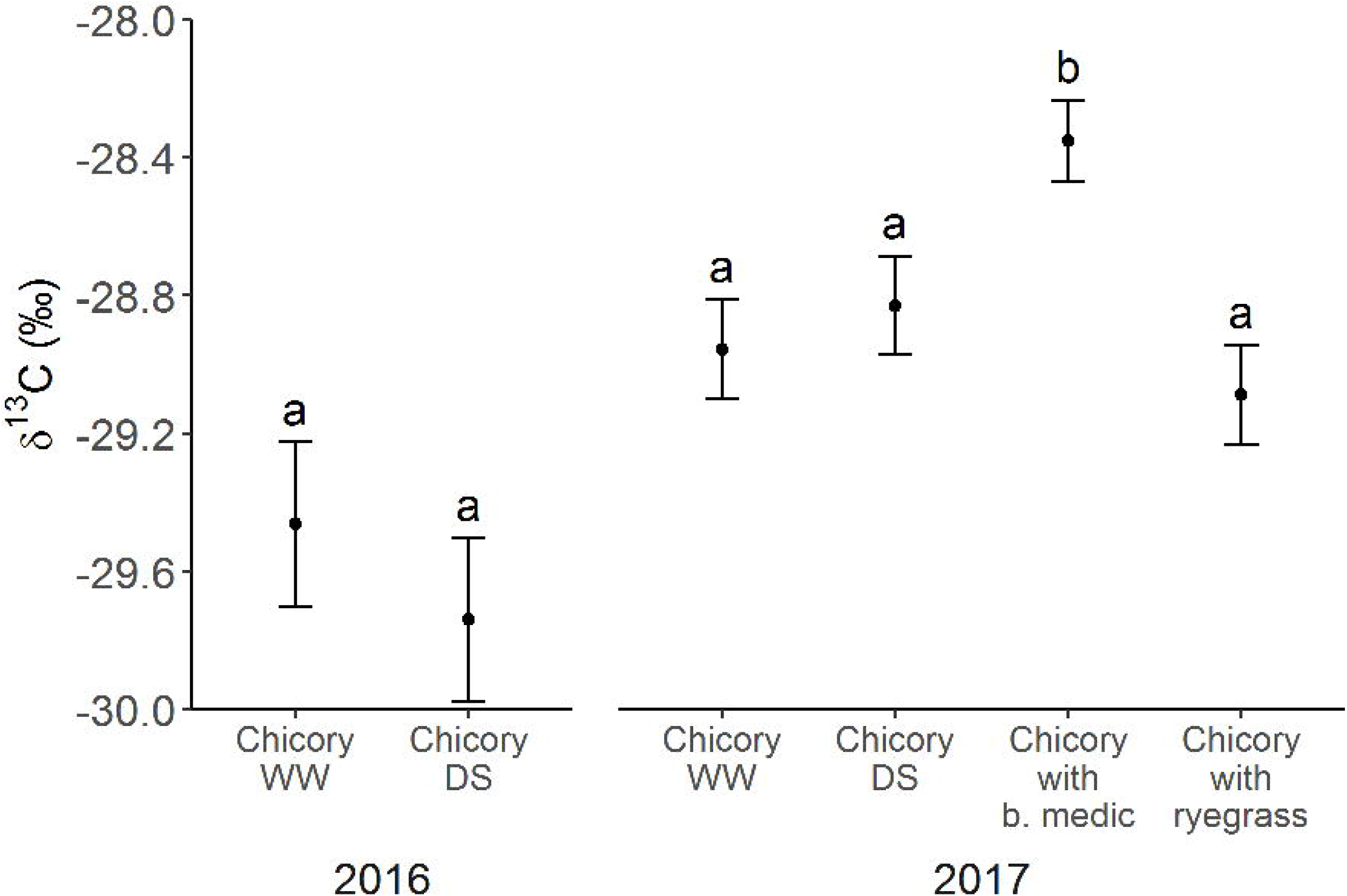
^13^C concentration in chicory harvested 28 July 2016 and 11 September 2017 in the well-watered (WW) and drought stressed (DS) chicory sole crop treatments and in the chicory intercropping treatment with ryegrass and black medic respectively. Error bars denote standard errors, and letters indicate significant differences among treatments in a one-way ANOVA. Separate analyses were made for each year. For 2017, the model is a mixed effects model because samples for each plant in a chamber were analysed separately

## Discussion

### Deep root growth

In accordance with our hypothesis, chicory demonstrated its capability to grow roots below 3 m depth and did so within 4.5 months. However, root intensity decreased markedly below 2 m in 2016 and below 2.5 m depth in 2017. The root intensity below 2 m depth at drought initiation, 2.5 m depth at tracer injection and 3.5 m depth at harvest in 2017 was very low and could be a result of roots from the 2016 crop still visible on the rhizotron surface. Studies covering a longer growing season have found extensive root growth in chicory down to 2.5 m, where equipment limitations prevented observations deeper down (Thorup-Kristensen 2006; Thorup-Kristensen and Rasmussen 2015). In the field, higher soil bulk density (Stirzaker et al. 1996; Gao et al. 2016) and other factors might restrict deep root growth, which is less likely in our semi-field facility with repacked soil. However, we did use field soil with a soil bulk density comparable to field soils.

Both intercropping treatments decreased total root intensity especially from 0.5 to 2 m depth. This has to be seen in the light of a total aboveground biomass that was twice as high as in the WW sole crop treatment. Observing that chicory biomass, on the other hand, was reduced to almost half when intercropped, suggests that both black medic and ryegrass had much lower root intensity below 0.3 m depth than sole cropped chicory and that the interspecific competition reduced both above- and belowground growth of chicory. Black medic and ryegrass are both shallow rooted and are unlikely to reach below 1 m depth (Kristensen and Thorup-Kristensen 2004; Thorup-Kristensen and Rasmussen 2015), thus the deep roots observed in the intercropping treatments are assumed to be chicory roots.

### Deep water uptake

The sensors documented water uptake in all treatments from 1.7 m depth in 2016 and 2.3 m depth in 2017. In fact, the sensors showed that in 2016, chicory water uptake at 1.7 m depth was c. 30 % of its water uptake at 0.5 m depth when well-watered. In 2017, chicory water uptake at 2.3 m depth was c. 10 % of its uptake at 0.5 m depth when well-watered. In absolute terms, water uptake from 1.7 m depth in 2016 was in the range of 1.5 mm m^-1^ soil column day^-1^ and from 2.3 m depth in 2017, it was 0.5 mm m^-1^ soil column day^-1^. Due to the small-sized plot placed at a windy position at 4 m height, evapotranspiration must have been substantially higher than the potential evapotranspiration measured nearby of 3.3 and 2.1 mm day^-1^ for the same periods in 2016 and 2017 respectively. Even though we did not estimate the total evapotranspiration, it is clear that the water uptake from the deeper part of the root zone substantially contributed to the total plant water balance.

The ^2^H tracer uptake by chicory from 2.3 m depth both years support the sensor-based water uptake calculations. Furthermore, the tracer study confirmed that neither black medic nor ryegrass had roots deep enough to acquire water from 2.3 m depth. This is a clear example of resource complementarity in root competition in intercropping (Tilman et al. 2001; Postma and Lynch 2012).

### Response to water stress and intercropping

Water uptake from 0.5 m depth was significantly reduced in the DS treatment compared to the WW treatment indicating that the soil water potential was low enough to limit plant water uptake in the DS treatment. Contrary to our expectations, we did not find a higher water uptake neither at 1.7 m depth in 2016 nor at 2.3 m depth in 2017 when plants were water limited in the topsoil. As biomass was not significantly reduced, whereas water uptake was reduced by 59 and 74 %, the reduction in water uptake cannot be explained by a reduced water need.

Although not significant, the ^2^H tracer study indicated a higher ^2^H concentration in the transpiration water in the DS compared to the WW treatment in 2016. This suggests a higher relative water uptake from 1.7 m depth. A higher relative uptake from a certain depth can logically be explained by an increase in water uptake from the given depth, a decrease in water uptake somewhere else in the soil profile or a combination of both. As the water uptake based on the sensor calculations show a significantly lower water uptake from 0.5 m depth in the DS than in the WW treatment in 2016, it is likely that what we observed was the effect of decreased uptake in the topsoil.

We only observed a significant increase in ^13^C concentration in chicory when intercropped with black medic. Samples were taken from the total biomass, and not from plant parts developed during the drought, which might explain why the treatment effects were only captured in the chicory and black medic intercropping, where black medic appeared to have induced drought stress in chicory even before the onset of the drought stress we induced.

Intercropping reduced total root intensity at 0.5 m depth by over 40 %. Still, water uptake from this depth was only slightly decreased indicating that the lower root intensity did not restrict water uptake in well-watered conditions. Root density in upper soil layers of well-established crops does not correlate well with water uptake (Anblin and Tennant 1987; Katayama et al. 2000), which can be explained by the high mobility of water in the soil, making a dense root system superfluous. Following the logic behind Walter’s two-layer hypothesis (Walter 1939, 1971; Walker and Noy-Meir 1982), intercropping would lead to a vertical niche partitioning resulting in increased water uptake by the deep-rooted chicory when intercropped with a shallow-rooted species. However, we did not observe an increase in deep water uptake.

### Absence of a deep water compensation effect

We suggest three possible explanations for why we did not observe the hypothesized increase in deep water uptake during drought or intercropping.

1) *The hydraulic resistance is too high to increase deep water uptake.* Theoretically, the ability of root systems to extract water from deep roots depends not only on root system depth but also on root system hydraulics (Javaux et al. 2013). Root hydraulic conductivity limits the potential water uptake, and differs among species, but also among different roots in a root system (Ahmed et al. 2018; Meunier et al. 2018). The ability of a root system to compensate, i.e. extract water where it is easily available, for instance from deeper soil depths, is, therefore, a function of (1) the xylem conductance between the roots in the extraction zone and the root crown and (2) the radial root conductance in the wet zone. Compensation has been observed in chicory below 0.6 m depth, but this was in a study allowing root growth down to only 1.5 m depth (Vandoorne et al. 2012a). In our experiment, the xylem conductance might simply have been too low in the deeper part of the root zone to allow compensation, possibly because the deep soil measurements were made in a zone with a low density of young roots (McCully 1995; Meunier et al. 2018). However, chicory had 31 % fewer roots in the chicory and black medic intercropping than in the WW treatment at 2.3 m, with no reduction in water uptake, not supporting such a relationship between root density and water uptake.

2) *Insufficient water supply in the topsoil induces root-to-shoot signalling causing stomata I closure, despite sufficient water supply in deeper soil layers.* Signals by phytohormones like Abscisic acid (ABA), produced when parts of the root system are under low water potential, might reduce plant transpiration and consequently root water uptake also from deeper depths by triggering stomatal closure (Zhang and Davies 1990a, b; Tardieu et al. 1992; Dodd et al. 2008). Split-root experiments, where one side of the root system is under low water potential, have found reduced stomata conductance, despite sufficient water supply (Blackman and Davies 1985; Zhang and Davies 1990b). However, experiments with vertical heterogeneity in soil water content yield ambiguous results (Puértolas et al. 2015; Saradadevi et al. 2016). The hormonal signalling during topsoil drying has not been tested for chicory. But chicory does show an isohydric behaviour, decreasing stomatal conductance and maintaining leaf water potential during moderate drought stress (Vandoorne et al. 2012b).

3) *Deep water uptake compensation might have occurred, but was not captured in this experimental set-up.* Water uptake compensation could have happened between or below the depths covered by the sensors. In 2016, VWC was not only lower at 0.5 m depth in the DS treatment compared to the WW treatment but also in 1.7 m depth, which could have impaired the water uptake from this depth, too. Water uptake could also have been confounded with water redistribution in the soil column, leading to an underestimation of water uptake in depths where water is moving to, and an overestimation in depths where water is moving from.

In summary, chicory can grow roots down to 3 m depth within 4.5 months and benefit from a significant water uptake from below 2 m depth both during well-watered and drought conditions. Our study highlights the benefit of deep root growth for crop water uptake, but questions whether further compensation in deep water uptake takes place when water is limited in the topsoil. A compensation might however, be pronounced for other crop species or for crops which have had more time to establish a deep root system.

## Acknowledgments

We thank statisticians Signe Marie Jensen and Helle Sørensen for advice regarding the statistical data analyses, technician Jason Allen Teem for his contribution to the experimental work, student helper Sunniva Melhus for counting roots and Engineer’s assistant Sebastian Nielsen for building the photo box and for constructive discussions on the design of it. All are affiliated with University of Copenhagen. We thank Villum Foundation (DeepFrontier project, grant number VKR023338) for financial support for this study.

